# Differential microstructural development within sensorimotor cortical regions: A diffusion MRI study in preterm and full-term infants

**DOI:** 10.1101/2025.06.07.658344

**Authors:** Alexandra Brandstaetter, Andrea Gondova, Laurie Devisscher, Denis Rivière, Guillaume Auzias, Yann Leprince, Jessica Dubois

## Abstract

The sensorimotor system develops early in utero and supports the emergence of body representations critical for perception, action, and interaction with environment. While somatotopic protomaps are already developed in the primary somatosensory and motor cortices in late pregnancy, little is known about the anatomical substrates of this functional specialization. In this study, we aimed to decipher the microstructural properties of these regions in the developing brain. Using advanced diffusion MRI and post-processing tools, we parcellated the pre- and post-central gyri into microstructurally distinct clusters along the lateral-to-medial axis in full-term neonates, confirming the early differentiation within sensorimotor regions. These clusters were further analyzed in preterm infants scanned near birth and at term-equivalent age (TEA), compared with another group of full-term neonates. Applying a multivariate Mahalanobis distance approach, we quantified deviations in preterm cortical microstructure relative to the full-term reference. Preterm infants showed significant region- and position-specific deviations at both ages, though these were smaller at TEA, consistently with ongoing maturation during the pre-term period. Differences between the pre- and post-central gyri, and along the somatotopic axis, suggested differential vulnerability to prematurity. In particular, the motor regions appeared to be at a more advanced stage of maturation close to birth and less vulnerable at TEA than somatosensory regions. An opposite trend was observed for lateral positions related to mouth representation compared with intermediary and medial positions. These findings support the notion that early sensorimotor cortical specialization is microstructurally emergent during gestation and sensitive to atypical developmental context of preterm birth.

## 1. Introduction

The sensorimotor system is one of the first to develop *in utero* in interaction with other sensory functions to facilitate the perception, integration and emotional processing of environmental information. In addition to early genetic and epigenetic factors, this early development is thought to depend on the multiple somatosensory stimulations and motor experiences of the foetus and then the baby (e.g., tactile contacts through the continuous pressure of the amniotic fluid over the body and with the mother’s uterine membrane, feedback from spontaneous motor behaviours) (de Vries et al., 1982; Fagard et al., 2018). At the neural level, the early establishment of thalamocortical afferent connections to the subplate (Kostovic et al., 2019) is supposed to provide the basis for the cortical processing of such sensorimotor activity from mid-gestation. Underlying cortical regions seem to be partly specialized at an early stage, as suggested by functional MRI (fMRI) studies of preterm and full-term neonates showing localized responses to tactile and passive motor stimulations (e.g., Allievi et al., 2016; Dall’Orso et al., 2018; Eyre et al., 2021). In particular, a proto somatotopy has been described during an equivalent period to the third trimester of gestation (Dall’Orso et al., 2018), with a topographic representation of body parts in the pre-central and post-central gyri (considered as primary somatosensory and motor cortices, respectively), from tongue to toes along the lateral-ventral to medial-dorsal axis of the classical ‘homunculus’ model reported in adults by (Penfield & Boldrey, 1937). A certain spatial organization of functional connections between these regions also seems to be present early on, leading to an adult-like configuration by the time of full-term birth (Dall’Orso et al., 2022). Such complex functional organization is assumed to be fundamental for motor skill acquisitions, imitation and social interactions during infancy (Marshall & Meltzoff, 2015).

Nevertheless, early developmental disturbances and alterations in environmental influences might impact such cerebral organization, with possible long-term consequences on the child behavioral acquisitions and learnings. A common example is preterm birth (i. e., delivery before 37 weeks of gestational age - w GA – which affects over 10% of newborns every year) (Martin et al., 2023). Preterm neonates experience strong changes in physiological and physical conditions (e.g., reduced movements due to gravity outside the womb) (Anderson, 1986; Fyfe et al., 2014; Teune et al., 2011), as well as early *ex utero* exposure to the environment of the neonatal intensive care unit (e.g., possible aggressive sounds, lights, limited tactile contacts), and painful medical procedures (Cong et al., 2017; Flacking et al., 2012). According to the embodied brain model, such exposure due to prematurity might impair the development of somatotopic organization (Yamada et al., 2016), though direct *in vivo* evidence is lacking.

Besides, a number of studies in preterm infants have shown developmental disturbances of the brain sensorimotor system, even in the absence of overt brain lesions, using advanced anatomical and diffusion MRI (dMRI) approaches (Dubois et al., 2021; Ouyang, Dubois, et al., 2019a). Notably, complex microstructural models (e.g., neurite orientation dispersion and density imaging NODDI) have appeared informative to analyse multi-shell dMRI data compared to the diffusion tensor imaging (DTI), and multiparametric methods may help integrating complementary measures. For example, our previous work demonstrated that white matter connections within the sensorimotor network are differentially impacted by prematurity at term equivalent age, with a caudal-to-rostral vulnerability that might mirror the typical progression of maturation (Neumane et al., 2022). Additionally, prematurity impacts the microstructural maturation of cortical regions across the whole brain (e.g. (Ball et al., 2013; Batalle et al., 2019; Gondova et al., 2025; Ouyang, Jeon, et al., 2019). Of note, over this early developmental period, advanced maturation of cortical grey matter is generally associated with a decrease in DTI-derived fractional anisotropy (FA), as well as axial, radial and mean diffusivities (AD, RD, MD) and an increase in NODDI-derived orientation dispersion index (ODI), while the pattern of neurite density index (NDI) is plausibly more complex (with a pre-term decrease followed by an increase), reflecting the growth of dendritic arborization and neurites, the increasing cellular and organelle density contrasting with cortical expansion (Batalle et al., 2019). In addition, prematurity seems to modulate the covariation of microstructural properties between cortical regions (named “microstructural connections”) differently in the sensorimotor network compared to the visual and associative networks (Gondova et al., 2025).

This underscores the importance of evaluating whether prematurity impacts the somatotopic organization of sensorimotor regions. While we might expect a global impact on the progression of specialization, we also hypothesize differential effects depending on which body parts become progressively functional and used. However, systematic mapping with fMRI in neonates remains challenging. As an alternative first step, we propose evaluating the microstructural properties of these cortical regions. Recent studies in adults have suggested strong links between body representations within the pre- and post-central gyri, the morphological patterns of the central sulcus separating them (Germann et al., 2020), and their myeloarchitecture (Kuehn et al., 2017). Furthermore, in a study comparing full-term neonates to adults, we showed that anatomically defined parcels of these gyri show distinct microstructural properties and a different progression of maturation over the perinatal weeks (Chauvel et al., 2020; Devisscher et al., 2021).

In this study, we then aimed to delineate regions within pre- and post-central gyri based on local microstructural properties at full-term birth and assess whether the resulting parcels are differentially affected by prematurity and whether they show different maturational patterns during a period equivalent to the last trimester of gestation. We used an advanced dMRI multi-parametric approach to compare preterm infants scanned close to birth and at term equivalent age with full-term neonates. We hypothesized that a data-driven method would allow us to reveal distinct cortical clusters within pre- and post-central gyri, which would have a somatotopic significance as suggested by mapping studies in adults. We further assumed that 1) sensory vs motor clusters, and 2) lateral vs medial clusters (related to different body parts) would show a different vulnerability to prematurity as well as a different developmental progression in the pre-term period, because of the asynchrony in behavioural acquisitions during this period.

## 2. Methods

### Data presentation

#### Subjects

This study included a sample of preterm and full-term neonates from the 3^rd^ release of the developing Human Connectome Project (dHCP) database (Edwards et al., 2022), collected at St Thomas’ Hospital, London, UK, between 2015 and 2020. The project received approval from the UK NHS Research Ethics Committee (14/LO/1169, IRAS 138070), and written informed consent was obtained from the parents of all participants.

Our analysis focused on four subject groups: PT_TEA_ consisting of 59 low-risk preterm-born infants scanned at term-equivalent age (TEA) (33 males and 26 females, median gestational age (GA) 31.7 weeks (w) [23.7w-36.0w]; median postmenstrual-age (PMA) at MRI 41.3w [38.4w-44.9w]); PT_Birth_ consisting of 45 subjects from the PT_TEA_ group that were scanned close to birth (26 males and 19 females, median GA at birth 32.3w [25.6w-36.0w]; median PMA at MRI 34.9w [28.3w-36.9w]). As a normative reference, we used a group of 59 full-term (FT) neonates matched to PT_TEA_ infants according to PMA at MRI and sex (FT_Ref_: median GA at birth 40.1w [37.4w-42.3w]; median PMA at MRI 41.3w [38.3w-44.3w]). Additionally, an independent group of 25 FT infants (FT_Ind_: median GA at birth 40.1w [37.1w-41.4w]; median PMA at MRI 41.3w [38.1w-43.9w]), with similar subject characteristics to FT_Ref_ in terms of PMA at MRI, GA at birth and sex distribution, was used for defining the cluster-based cortical parcellation. Subject characteristics are detailed in *Table 1*. See *Supp. Fig. 1* for distribution of ages for the four cohorts.

**Table 1.** Group characteristics. *Legend*: FT_Ind_: independent full-term group, FT_Ref_: reference full-term group, GA: gestational age, N: number, PMA: post-menstrual age, PT_Birth_: preterm group scanned close to birth, PT_TEA_: preterm group scanned at term equivalent age (TEA).

|  | PT<sub>Birth</sub> | PT<sub>TEA</sub> | FT<sub>Ref</sub> | FT<sub>Ind</sub> |
| --- | --- | --- | --- | --- |
| N | 45 | 59 | 59 | 25 |
| Sex: |  |  |  |  |
| males | 26 | 33 | 33 | 15 |

| females | 19 | 26 | 26 | 10 |
| --- | --- | --- | --- | --- |
| Median GA at Birth [range]<br>in weeks | 32.3 [25.6 – 36.0] | 31.7 [23.7-36.0] | 40.1 [37.1 – 42.3] | 40.1 [37.1 – 41.1] |
| Median PMA at Scan [range]<br>in weeks | 34.9 [28.3 – 36.9] | 41.3 [38.4 – 44.9] | 41.3 [38.3 – 44.3] | 41.3 [38.3 – 43.9] |

**Figure 1.**
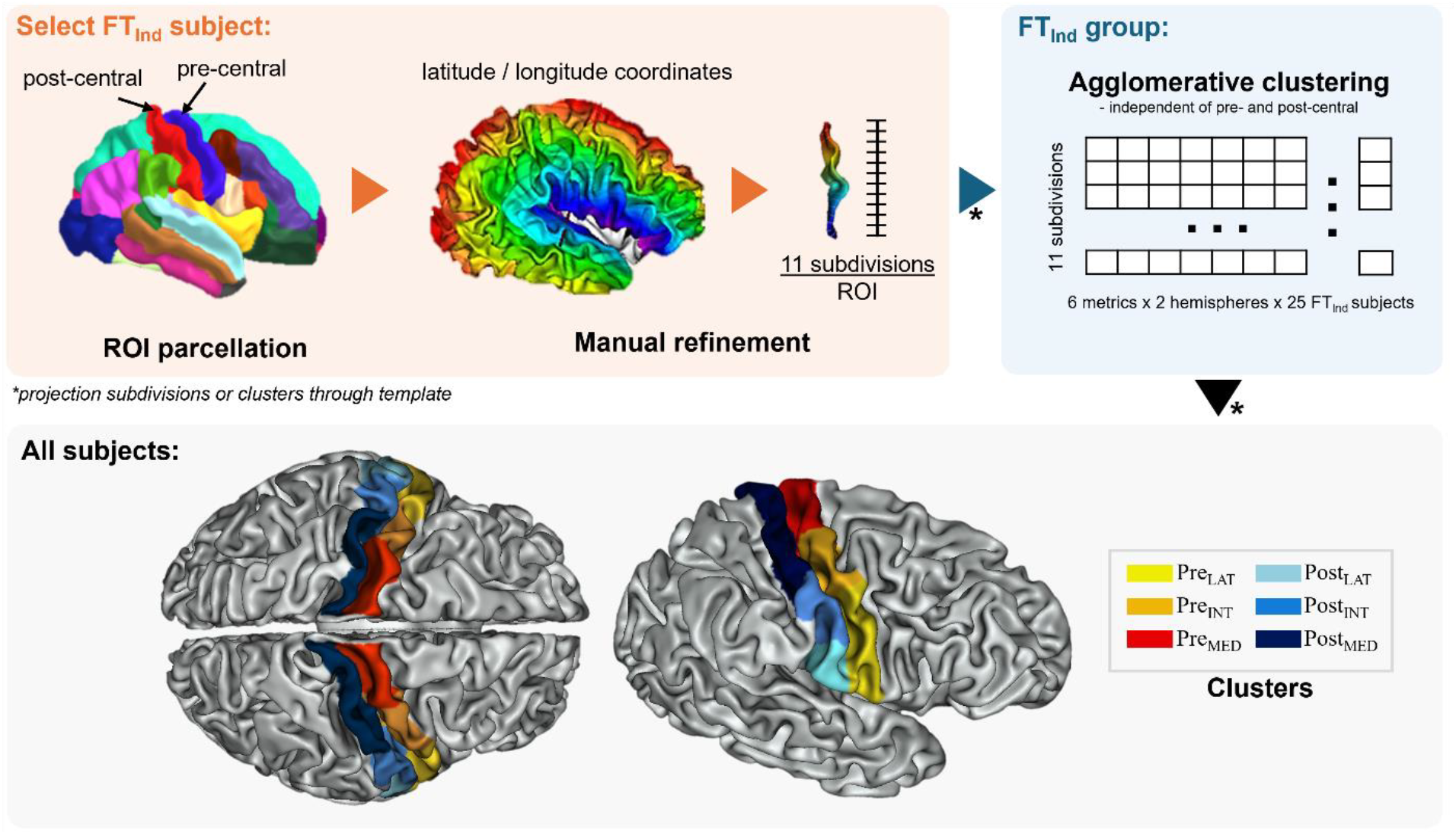
Summary of the pipeline for microstructure-based clustering of pre- and post-central regions of interest (ROI) used in our work as well as visualization of the resulting clusters. *Legend:* Pre_LAT_ / Post_LAT_: lateral portion of pre-central gyrus (yellow) / lateral portion of post-central gyrus (light blue), Pre_INT_ / Post_INT_: intermediary portion of pre-central gyrus (orange) / intermediary portion of post-central gyrus (blue), Pre_MED_ / Post_MED_: medial portion of pre-central gyrus (red) / medial portion of post-central gyrus (indigo).

### Acquisition and preprocessing of MRI data

MRI data were acquired using a Philips 3 Tesla Achieva scanner (Philips Medical Systems, Best, Netherlands). All infants were scanned during natural sleep using a neonatal head coil and imaging system optimized for the dHCP study as previously described (Hughes et al., 2017). Here, we considered anatomical and diffusion MRI data available in its pre-processed state (Edwards et al., 2022).

The *anatomical data* resulted from acquisition and reconstruction of T2w images using optimized protocols (Cordero-Grande et al., 2019), leading to super-resolved images with an isotropic spatial voxel size of 0.5 mm. Processing followed a dedicated pipeline for brain tissue segmentation and cortical surface extraction of neonatal brains (Makropoulos et al., 2018), with bias-correction, brain extraction, volumetric segmentation using Draw-EM (Developing brain Region Annotation With Expectation Maximization) algorithm (Makropoulos et al., 2014). Interface between cortex and white matter (*wm surfaces*) were then extracted following an adapted pipeline with Morphologist toolbox (BrainVISA 5.1.1) (Rivière et al., 2009).

*Diffusion MRI data* (dMRI) were acquired following a multi-shell high angular resolution diffusion imaging (HARDI) protocol with 4 b-shells (b = 0 s/mm2: 20 repeats; and b = 400, 1,000, 2,600 s/mm2: 64, 88, and 128 directions, respectively) (Hutter et al., 2018) and pre-processed with correction for motion artifacts and slice-to-volume reconstruction using the SHARD approach, leading to an isotropic voxel size of 1.5 mm (Christiaens et al., 2021).

### Diffusion modelling and metric extraction

For each subject, the DTI model was fitted to the diffusion data using a single shell (b = 1,000 s/mm2) and calculated with FSL’s DTIFIT to estimate metric maps for 4 metrics: axial diffusivity (AD), radial diffusivity (RD), and mean diffusivity (MD), fractional anisotropy (FA). Additionally, multi-shell diffusion data was used to derive the neurite density index (NDI) and orientation dispersion index (ODI) maps from the NODDI model (Zhang et al., 2012) using the CUDA 9.1 Diffusion Modelling Toolbox (cuDIMOT) NODDI Watson model implementation for GPUs (Hernandez-Fernandez et al., 2019). Derived NODDI maps were then corrected as described in (Neumane et al., 2022). Diffusion metrics were then computed within the cortical ribbon through projection to the *wm surface* using a cylindrical approach guided by the minimum of AD, as described in (Gondova et al., 2023; Lebenberg et al., 2019).

### Microstructure-based clustering of pre- and post-central gyri

To derive microstructure-based clusters of the pre- and post-central cortical gyri distributed along the latero-medial axis in order to investigate their putative correspondence with somatotopic mapping, we first selected a single subject from the FT_Ind_ group and the two gyri were delineated bilaterally using the M-CRIB-S surface-based parcellation tool optimized for term-born neonates (Adamson et al., 2020). We then subdivided each of the four parcels (2 gyri x 2 hemispheres) into 11 subdivisions providing a sufficient degree of details and of roughly similar size. This was guided by *geometrical coordinates* along the central sulcus reconstructed using an adapted Sulcus Parametrization toolbox (Cykowski et al., 2008; Dornier et al., 2025), and *latitude/longitude coordinates* along the gyri computed with Hip-Hop Cortical Parameterization of the Cortical Surface (Auzias et al., 2013) (both toolboxes available in BrainVISA 5.1.1). To refine region definitions and reduce the number of regions, we aimed to further cluster the initial subdivisions based on their microstructural similarity. We first projected the 11 subdivisions per region into the anatomical space of the remaining FT_Ind_ subjects using non-linear transformations via the 40w dHCP template from the dHCP database that were computed using Multimodal Surface Matching (MSM) approach (Bozek et al., 2018). For each subdivision at the subject level, we extracted the median values of the 6 diffusion metrics, then concatenated these across FT_Ind_ subjects and hemispheres (to ensure correspondence between hemispheres during clustering) resulting in a gyrus-specific vector of length 132 (11 subdivisions x 2 hemispheres x 6 metrics). We then applied agglomerative hierarchical clustering with average linkage and Euclidean distance independently for the pre- and post-central gyri. The final number of clusters was selected based on visual inspection of the dendrograms and spatial coherence across hemispheres. These clusters were then projected to all infants of the PT_Birth_, PT_TEA_, and FT_Ref_ groups as described earlier. Cluster borders were manually refined, and DTI and NODDI metrics were extracted for each cluster and subject. Summary of the pipeline is visualised in *Figure 1*.

To assess the validity our microstructure-based clusters, we conducted t-tests for each median metric per cluster to ensure the two FT groups did not significantly differ in terms of microstructure within each cluster. Throughout our work, multiple comparisons were corrected using the Benjamini-Hochberg False Discovery Rate (FDR) method, with α=0.05 for all tests. Additionally, we performed a repeated-measure analysis of covariance (ANCOVA) across the PT_TEA_ and FT_Ref_ groups to provide a first evaluation of the effects of prematurity (group effect), and clusters (region effect: pre-vs post-central; position effect: lateral, intermediary vs medial) on diffusion metrics. Covariates included (1) PMA at MRI, and (2) residual whole-cortex metrics corrected for PMA (i.e. residuals from a linear regression of whole-cortex metric with PMA) to account for inter-subject variability in cortical maturation. The hierarchical structure of the repeated measures (matched PT-FT subjects across the two regions and three positions) were accounted for using a nested error term. For all analyses, statistical implementations were performed using R (version 2022.07.1) and the Pingouin (v0.5.4) Python package.

### Multivariate analysis of prematurity and maturation effects within pre- and post-central gyri

To further characterise the impact of prematurity and maturation on the microstructural profiles of PT subjects compared to FT_Ref_ subjects for each cortical cluster, we applied a multivariate approach based on Mahalanobis distance accounting for the 6 diffusion metrics. This method measures the distance between an individual PT subject and the FT_Ref_ group as a reference, accounting for inter-subject variability within the FT_Ref_ group and the collinearity between diffusion metrics. A smaller distance indicates that the PT subject is closer to the reference group. This approach has previously been successfully used to study differential maturation across white matter tracts during infancy (Kulikova et al., 2015) and the effects of prematurity across sensorimotor tracts (Neumane et al., 2022).

To compute the Mahalanobis distances, we first calculated the median of each diffusion metric (AD, RD, MD, FA, NDI, ODI) for each cluster across the two hemispheres, scaled the values to a [0; 1] range across the clusters for each metric and group, and corrected them within each subject group for PMA at can, and for corresponding residual diffusion metrics over the whole cortex (corrected for PMA at scan). We then computed the Mahalanobis distance in each cluster for both PT groups, using FT_Ref_ infants as the reference group representing typical variation of cluster brain microstructural characteristics at TEA. The Mahalanobis distance in PT_TEA_ infants (N = 59) would reflect the cluster’s deviation due to prematurity, while in PT_Birth_ infants (N = 45), the distance would indicate the level of immaturity during the preterm period.

We assessed whether PT groups differed from FT_Ref_ group by testing the nullity of the Mahalanobis distance using t-tests for all clusters corrected for multiple comparisons. We also conducted additional ANOVA analyses within each PT group to quantify the effects of different regions (pre/post-central) and positions (from lateral to medial) on the estimated Mahalanobis distance. We then examined the relationship between the Mahalanobis distances and PT subjects’ GA at birth using linear regression to assess whether the level of prematurity would relate to the clusters’ degree of maturation.

## 3. Results

### Microstructure-based clusters of the pre- and post-central gyri

Visual inspection of dendrograms resulting from the clustering analysis of the 11 subdivisions across the FT_Ind_ group revealed clear clusters in both the pre-central and post-central gyri *(Supp. Figure 2)*. Three homotopic clusters were identified in each gyrus of the left and right hemispheres (*Figure 1*), along the lateral-to-intermediary-to-medial position of both gyri: Pre_LAT_, Pre_INT_, Pre_MED_ for the pre-central gyrus, and Post_LAT_, Post_INT_, Post_MED_ for the post-central gyrus. As expected, the two FT groups did not show significant differences across the median diffusion metrics per cluster, as assessed using t-tests, except for ODI (*Supp. Table 1*), suggesting similar microstructural characteristics and equivalent representation of normal brain development across computational steps in our study.

When the identified clusters were considered to characterize the microstructural profiles of the groups of interest (PT_Birth_, PT_TEA_, and FT_Ref_), we observed variations across pre-central and post-central regions, positions and groups (*Figure 2*). Visual observations suggested a consistent spatial pattern along the lateral-to-medial axis in both gyri (e.g., higher diffusivity values in lateral than medial clusters). And as expected, systematic differences between the PT_Birth_ group compared to the PT_TEA_ and FT_Ref_ groups were visible: higher diffusivity and FA values contrasting with lower NDI and ODI values in PT_Birth_ infants, although ODI showed considerable variability within this group.

**Figure 2.**
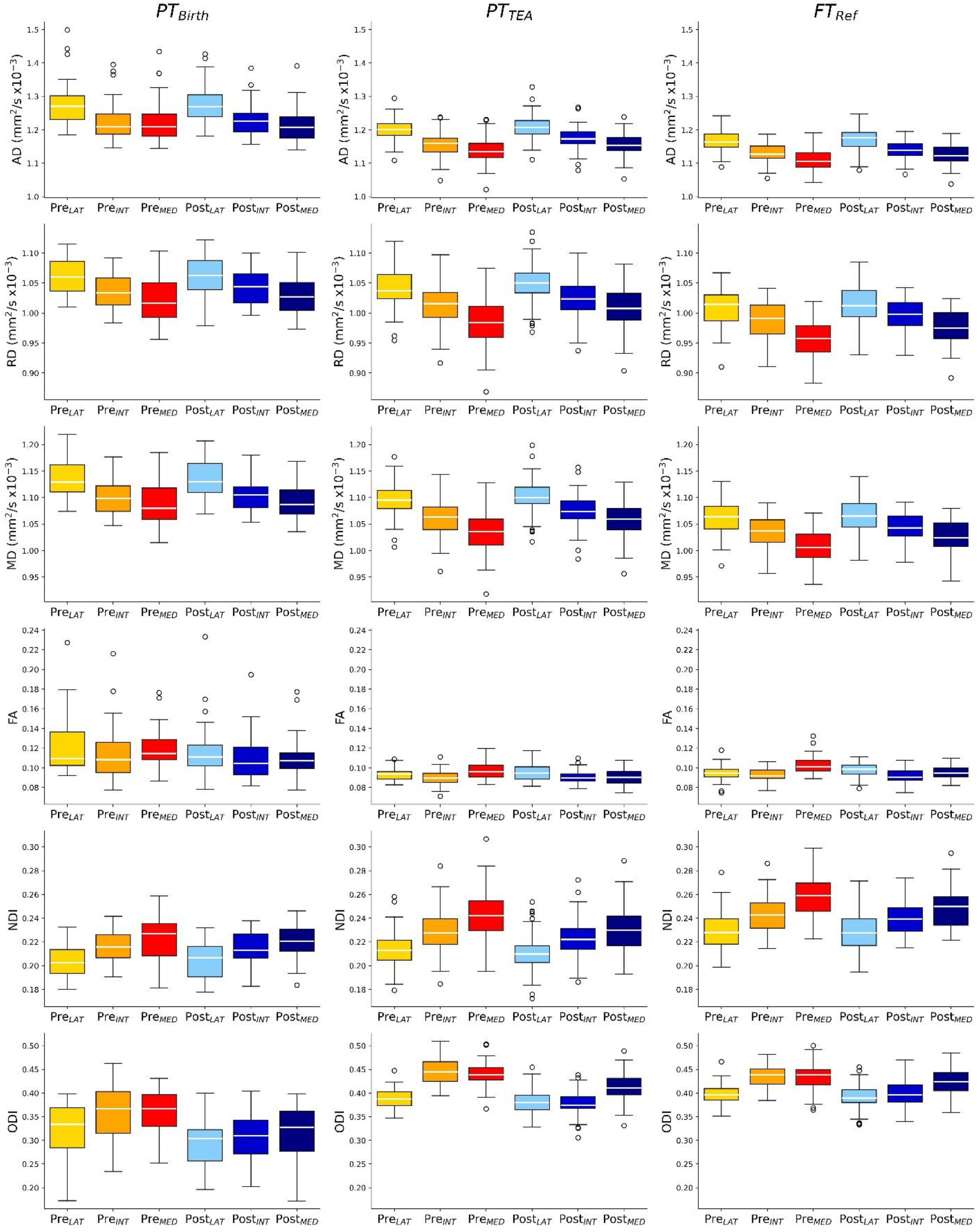
Diffusion metrics across subject groups and pre- and post-central clusters. *Legend*: AD: axial diffusivity, RD: radial diffusivity, MD: mean diffusivity, FA: fractional anisotropy, NDI: neurite density index, ODI: orientation dispersion index. See Table 1 for subject group legend and Figure 1 for cluster legend and colour-codes.

Further comparisons of the PT_TEA_ and FT_Ref_ groups using ANCOVA modelling on each individual diffusion metric confirmed the influence of prematurity on microstructural characteristics (group effect for all metrics), as well as significant effects of region and position *(Supp. Table 2)*.

Taken together, these first results highlighted the importance of analysing clusters separately within each region and position, as they showed distinct effects on diffusion metrics, suggesting variations in underlying microstructural properties and potentially different impact of prematurity and maturation. Nevertheless, the variability across individual metrics complicated the identification of clear patterns. A multivariate framework was thus needed to better capture and interpret prematurity effects and maturational differences across clusters, as outlined below.

### Multivariate analysis of the impact of prematurity and maturation on microstructural profiles of pre- and post-central gyri

Mahalanobis distances for the 6 bilateral clusters for the two preterm groups are summarized in Figure 3a, with lower values indicating that a subject’s microstructural profile is more similar to the full-term reference (FT_Ref_) distribution. By examining the Mahalanobis distances in the PT_Birth_ group, we can infer the degree of maturational changes within each cluster during the preterm period. In this group, all clusters showed significant deviations from 0 (*Supp. Table 3*), suggesting detectable ongoing maturational changes. The ANCOVA analysis over this group showed significant effects of both region (pre- and post-central gyri) and position, indicating heterogeneous microstructural maturation across the two gyri and the lateral to medial axis, but no effect of gestational age at birth. Pre_MED_ showed the smallest Mahanobis distance, suggesting it might be the most developmentally advanced among the clusters close to birth. In contrast, Pre_INT_ and Pre_LAT_ displayed larger distances indicating they will undergo more pronounced maturational changes during the preterm period. A similar gradient somewhat appears to be present for the clusters in the post-central gyrus. Post-hoc tests, conducted after concatenating values across positions for this group, confirmed significantly lower Mahalanobis distances in the pre-central gyrus compared to the post-central gyrus, suggesting more advanced maturation (paired t-test: T = -4.388, p < 0.001). Higher Mahalanobis distances were observed in the lateral clusters when values were concatenated across regions, suggesting less advanced maturation (LAT vs. INT: T = -5.933, p < 0.001; LAT vs. MED: T = -4.712, p < 0.001), with no significant differences between INT and MED positions (T = -0.380, p = 0.705).

**Figure 3.**
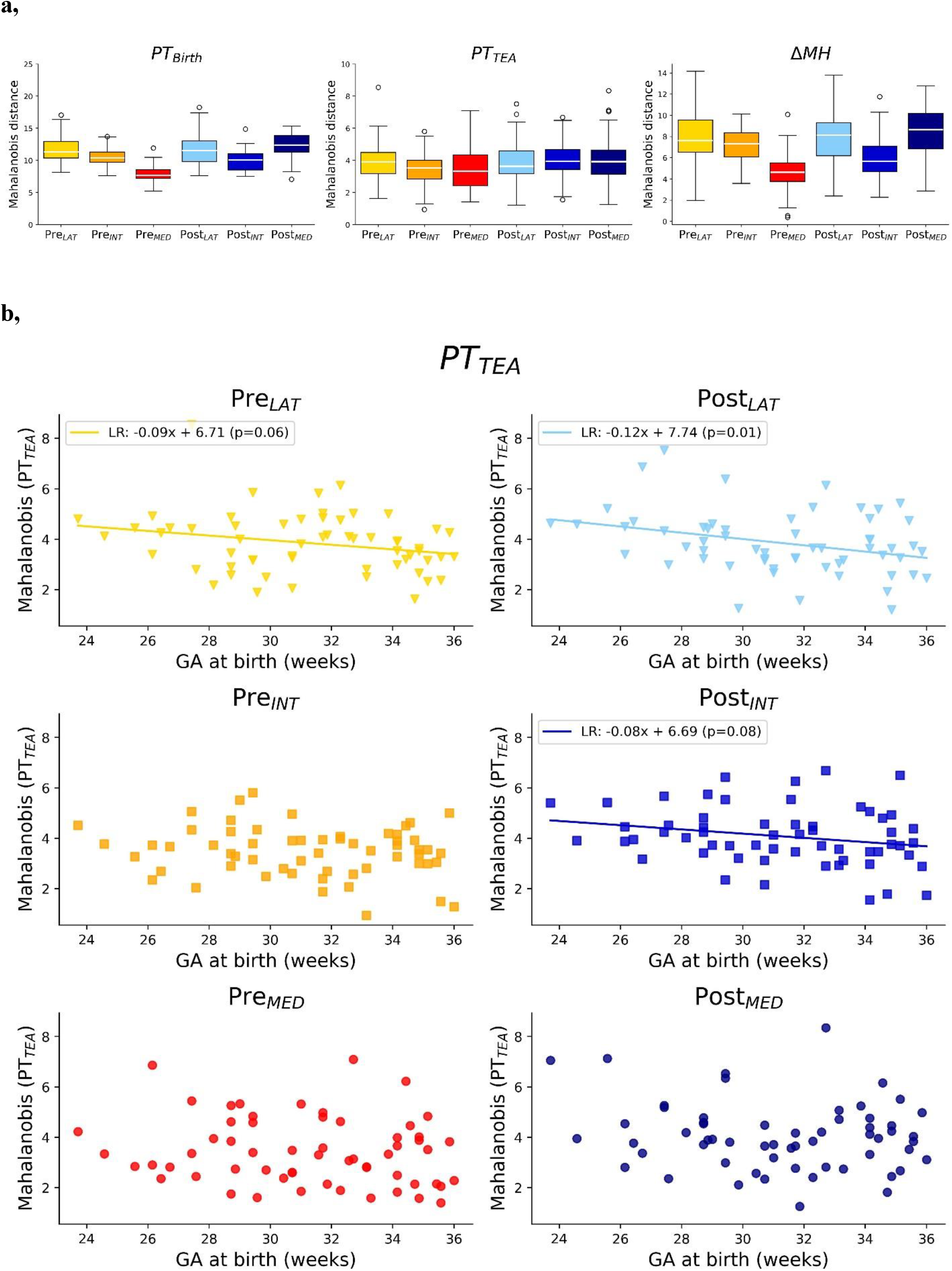
**a.** Mahalanobis distances across clusters for PT_Birth_ (left), PT_TEA_ (middle) groups, as well as Mahalanobis distance differences between the two sessions for the subjects with longitudinal data. **b**, Scatter plots presenting relationship of Mahalanobis distance at TEA with GA at birth (LR: line of linear regression with p<0.01). See Table 1 for subject group legend and Figure 1 for cluster legend and colour-codes.

In the PT_TEA_ group, assessing the Mahalanobis distances provides insight into the heterogenous effects of prematurity on microstructural development across the clusters at term equivalent age. Although the distances were substantially lower than those observed in the PT_Birth_ group (Figure 3a) —suggesting that microstructural maturation continued *ex utero* and began to converge toward the profile of full-term (FT) infants—all regions and positions exhibited Mahalanobis distances significantly different from zero (*Supp. Table 3*). ANCOVA modelling over this group highlighted a significant main effect of region, indicating that prematurity differentially affected the sensory and motor regions, but no main effect of position (*Table 2*). Although the patterns for the effect of region were less easy to discern visually in the PT_TEA_ group, post-hoc test confirmed lower distances in the pre-vs post-central gyrus (paired t-test: T=-4.169, p<0.001), in a similar way to the PT_Birth_ group. The ANCOVA in the PT_TEA_ group also showed a significant effect of GA at birth, that we investigated further using linear regression. Although visually weak negative slopes were observed across all clusters —suggesting that higher GA at birth is associated with microstructure closer to full-term norms at TEA— this relationship reached statistical significance (α = 0.05) only for the lateral post-central cluster (Post_LAT_) but trends for Post_INT_ and Pre_LAT_ (*Figure 3b*).

**Table 2.** For each PT group, ANCOVA analysis examining the relationship between Mahalanobis distance and GA at scan together with key factors: region (pre-/post-central gyrus), and position and their interaction. The models account for hierarchical structure of the repeated measures across matched subjects, regions, and positions, with a nested error term.

|  | MH: PT<sub>Birth</sub> |  |  | MH: PT<sub>TEA</sub> |  |  |
| --- | --- | --- | --- | --- | --- | --- |
| | F | p | Partial $\eta^2$ | F | p | Partial $\eta^2$ |
| *Covariates* |  |  |  |  |  |  |
| GA at birth | 0.25 | 0.619 | 0.006 | 6.086 | 0.017 | 0.096 |
| *Factors* |  |  |  |  |  |  |
| Region | 25.48 | <0.001 | 0.367 | 14.87 | <0.001 | 0.204 |
| Position | 16.06 | <0.001 | 0.157 | 0.356 | 0.701 | 0.003 |

## 4. Discussion

In this study of human infants, we parcellated the pre- and post-central gyri each into three cortical clusters along the lateral-ventral to medial-dorsal axis using diffusion MRI metrics. These clusters showed distinct microstructural profiles in both full-term neonates and preterm infants, including those scanned at TEA and close to birth, despite the effect of prematurity. Using a multivariate approach (Mahalanobis distance), we confirmed that preterm infants deviated from full-term reference at both time points, though deviations were smaller at TEA, consistent with ongoing maturation. Notably, the extent and pattern of deviations differed between the pre- and post-central gyri at both ages, suggesting differential effects of prematurity and maturation in motor versus sensory regions. Additionally, regional differences were also observed along the lateral-to-medial axis close to birth, supporting the hypothesis of asynchronous microstructural development across regions that become specialized for different body parts during the third trimester.

### Parcellating cortical regions based on microstructural features

Cortical microstructure was assessed with a dedicated approach using advanced diffusion MRI and benefitting from the complementary information provided by DTI and NODDI models. To ensure reliable quantification within the thin cortical ribbon of infants, we used an optimized methodology for projecting diffusion metrics onto the cortical surface that limits the partial volume effects from adjacent white matter and cerebrospinal fluid (Gondova et al., 2023; Lebenberg et al., 2019).

Previous studies examining similar developmental windows have reported a systematic decrease in DTI diffusivity metrics with advancing cortical maturation (Ball et al., 2013; McKinstry et al., 2002; Ouyang, Dubois, et al., 2019b), supposed to reflect reduction in water content due to increasing cellular and organelle density. Correspondingly, one might expect an increase in NDI, although recent findings have been less consistent in this regard (Batalle et al., 2019). Besides, the maturation-related increase in dendritic arbor complexity (Huttenlocher & de Courten, 1987) and the decreasing radial organization of the cortex (Kostovic et al., 2019) may account for observed decreases in DTI anisotropy and ODI reported during the preterm period.

Our findings align with these reported developmental trends related to cortical maturation. We observed the highest diffusivity and FA values, along with the lowest NDI and ODI, in preterms close to birth. These metrics showed intermediate values in preterms at TEA, and most mature profiles in full-term neonates. Among the two full-term groups, dMRI metrics were largely similar, except for ODI. This difference may reflect methodological factors, such as the potentially suboptimal parametrization of the NODDI model for neonatal brain tissue or variable grey-white matter segmentation quality across subjects – particularly relevant given ODI’s sensitivity to local tissue complexity like neurite orientation and dispersion that other metrics are less able to capture. Alternatively, group variability in GA at birth and/or PMA at scan and non-linear developmental changes around the perinatal period (Batalle et al., 2019) might have contributed to this ODI difference between the two full-term groups.

To further investigate regional microstructural variation within pre- and post-central gyri, we used an exploratory, data-driven clustering approach on dMRI metrics from an independent group of full-term neonates. Six diffusion-derived metrics were included, based on evidence that multimodal approaches improve the accuracy of microstructural evaluations compared to unimodal methods (Kulikova et al., 2015; Vaher et al., 2022). Although inter-metric correlations were not explicitly accounted for in our modelling, the spatial consistency of the resulting clusters supports the validity of the method. Our approach combined initial manual delineation into 11 subdivisions per region with agglomerative clustering to refine and reduce cluster numbers. While direct vertex-wise clustering could offer a more purely data-driven solution, it might produce fragmented, spatially inconsistent regions. The two-step method thus balances anatomical plausibility with data-driven refinement. For simplicity and statistical power, left and right hemispheres were analysed together, which precluded assessment of inter-hemispheric asymmetries — current limitation that may be addressed in future studies.

Additionally, while the cortical clusters were defined based on full-term data, they were subsequently used to compare preterm infants to full-term references. Clustering each group independently would have impeded cross-group comparisons. Although including preterm infants in the clustering phase might have mitigated this issue, it was not possible due to limited availability of independent preterm data. Thus, a potential limitation of our approach is that regions defined on full-term brains may not perfectly correspond to preterm brain anatomy—differences in region size or position could cause some overlap with neighbouring areas and signal contamination. However, preterm signals—especially at birth—exhibit more extreme rather than blended values, suggesting that mapping inaccuracies are unlikely to drive the observed group differences. Moreover, the consistent microstructural patterns observed across all groups in diffusion metrics support the robustness of the cluster definitions and justify their use in comparative analyses.

### Characterizing microstructural profiles of the pre- and post-central gyri

We parcellated the pre- and post-central gyri along the lateral-to-medial axis into spatially coherent and consistent clusters in full-term neonates. An ANCOVA on dMRI metrics across groups scanned at TEA confirmed distinct microstructural profiles across the clusters (main effect of position) and also suggested systematic differences between the pre- and post-central gyri (main effect of region). Similar microstructural differences have already been reported in adult brain, with studies showing particularly a higher orientation dispersion and myelination in the pre-compared to post-central gyrus, based on dMRI metrics (Fukutomi et al., 2018), and T1w/T2w ratio-derived myelination metric (Kuehn et al., 2017).

Our findings suggest that such regional differences in microstructure might emerge early in cortical development. Despite variability in PMA at scan – and associated differences in brain size and gyrification – preterm neonates at both time points also showed consistent patterns of differences in diffusion metrics across both regions and along the lateral-to-medial axis. However, we refrained from directly interpreting these spatial differences in terms of maturation due to the absence of normative, mature reference values for neonatal dMRI metrics. This would have required comparisons with adult datasets (Dubois et al., 2016), to enable the computation of maturational distances (Devisscher et al., 2021; Kulikova et al., 2015). Yet, such comparisons would be methodologically challenging, given differences in brain size, cortical folding, and diffusion acquisition protocols across age groups.

Additionally, we remain cautious interpreting the potential functional significance of the identified clusters, as somatotopic mapping data were not available for the dHCP cohort. This limits our ability to interpret clusters in terms of specific body part representations. In adults, the lateral pre- and post-central regions typically correspond to facial areas, intermediate regions to the hands, and medial regions to the lower limbs. A visual comparison between our clusters and the only publicly available infant fMRI somatotopic maps (mouth, wrists, ankles; Dall’Orso et al., 2018) revealed neither a clear correspondence nor an outright mismatch. Thus, the relationship between early cortical microstructure and emerging functional specialization remains an open question.

### Highlighting the effects of prematurity and maturation on sensorimotor regions

To better capture differences between infant groups, we adopted a multivariate approach to compare preterm infants to a full-term reference, as proposed in previous study of sensorimotor white matter pathways in preterm infants at TEA (Neumane et al., 2022). Multivariate models, by leveraging inter-metric complementarity and typical individual variability, can more effectively capture complex, subtle group differences compared to univariate analyses. Here, we computed Mahalanobis distances using scaled dMRI metrics, with correction for PMA at scan and metric residuals over the whole cortex. Gestational age (GA) at birth was not included as a covariate, as it had minimal influence on individual metrics (data not shown) and was instead a variable of interest to evaluate whether GA influenced the prematurity distances. Sex was also not controlled for due to limited sample size, although it may contribute to inter-individual variability.

Using this approach, we found that preterm infants, both near birth and at term-equivalent age (TEA), significantly deviated from full-term profiles. These deviations were smaller at TEA, consistent with ongoing maturation. Our findings align with recent work showing region-specific cortical maturation trajectories (Ball et al., 2013; Batalle et al., 2019; Ouyang, Dubois, et al., 2019b), as well as heterogeneous regional effects of prematurity on cortical microstructure (Gondova et al., 2025; Ouyang, Jeon, et al., 2019). Our study extended these observations by providing a more fine-grained analysis within the sensorimotor cortex. Close to birth, significant main effects of both region (pre-vs post-central gyrus) and position (lateral to medial) were found. The pre-central gyrus exhibited lesser impact of prematurity than the post-central gyrus, and lateral clusters appeared less impacted than intermediate and medial ones. Interestingly, medial clusters displayed divergent microstructural profiles across the pre- and post-central gyri compared to other positions, which may be attributed to unequal sizes of medial clusters, with Post_MED_ cluster spanning a relatively large portion of the cortex. At TEA, only the main effect of region (pre-vs post-central) remained significant, though the positional trend persisted within the pre-central gyrus.

Patterns across our results may be interpreted through two organizing principles. First, motor cortices appear to be more mature than sensory cortices during the preterm period and less affected by prematurity at TEA. This is consistent with our previous findings in white matter pathways, where motor connections were less vulnerable to prematurity than sensory pathways (Neumane et al., 2022). Second, cortical regions corresponding to the mouth showed more pronounced immaturity and greater sensitivity to prematurity than those associated with the hands or lower limbs. This was somewhat unexpected given the early and intense functional use of the mouth in fetuses and neonates (e.g., intense suckling perceptions and experiences). Traditional developmental models suggest functional progression from top to bottom, and from medial to distal regions; however, our cortical findings align more closely with established neurodevelopmental gradients—particularly a caudal-to-rostral, and possibly medial-to-lateral pattern (Dubois et al., 2014; Yakovlev & Lecours, 1967).

Moreover, in approximately half of the cortical clusters, the Mahalanobis distance between preterm infants at TEA and full-terms systematically decreased with increasing GA at birth. This finding highlights that the degree of prematurity (extreme, very, moderate, late) significantly influences cortical microstructure, even at term-equivalent age. Collectively, our results support the notion that regions maturing later are more susceptible to prematurity-related disruption, emphasizing that the timing of preterm birth relative to a region’s developmental trajectory is a critical determinant of its vulnerability.

## 5. Conclusion

Our findings demonstrate that the microstructural organization of the pre-central and post-central gyri is both region- and position-specific, emerges early in development, and is differentially influenced by prematurity. The absence of neonatal functional data limits the interpretation of these results, particularly since structurally immature regions—such as the mouth area—are known to be functionally active early on, highlighting a potential dissociation between structural maturation and functional engagement. Future research, especially longitudinal studies integrating microstructure, function, and behaviour in preterm infants, will be essential to elucidate how these early structural differences relate to functional specialization and long-term neurodevelopmental outcomes.

## CRediT authorship contribution statement

**Alexandra Brandstaetter:** Writing – original draft, Writing – review & editing, Methodology, Formal analysis, Data curation. **Andrea Gondová:** Writing – original draft, Writing – review & editing, Methodology, Formal analysis, Validation, Visualization. **Laurie Devisscher-Gheude:** Writing – review & editing, Methodology. **Denis Rivière:** Writing – review & editing, Methodology. **Guillaume Auzias:** Writing – review & editing, Methodology. **Yann Leprince:** Writing – review & editing, Methodology. **Jessica Dubois:** Writing – original draft, Writing – review & editing, Methodology, Validation, Supervision, Funding acquisition, Conceptualization.

## Acknowledgments

We thank Prof. Alard Roebroeck of the Faculty for Psychology and Neuroscience at Maastricht University, who provided valuable support and guidance throughout the course of the Master’s thesis project that forms the basis of this work.

