## Supplementary Materials for "Differential microstructural development within sensorimotor cortical regions: A diffusion MRI study in preterm and full-term infants"

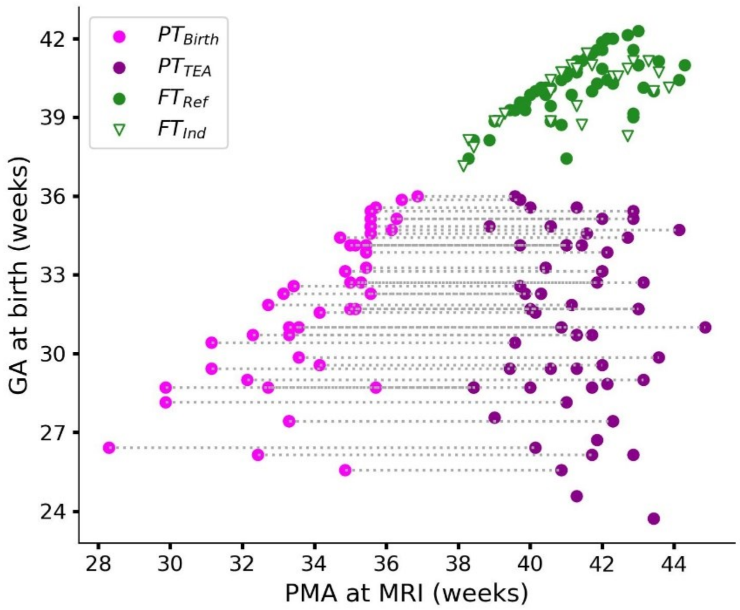


**Supp. Figure 1.** Distribution of post-menstrual age (PMA) at MRI and gestational age (GA) at birth for the 4 subject groups. Grey dotted lines connect longitudinal scans for the same subject. *Legend*: $PT_{Birth}$: preterm group scanned close to birth, $PT_{TEA}$: preterm group scanned at term equivalent age (TEA), $FT_{Ref}$: reference full-term group, $FT_{Ind}$: independent full-term group.

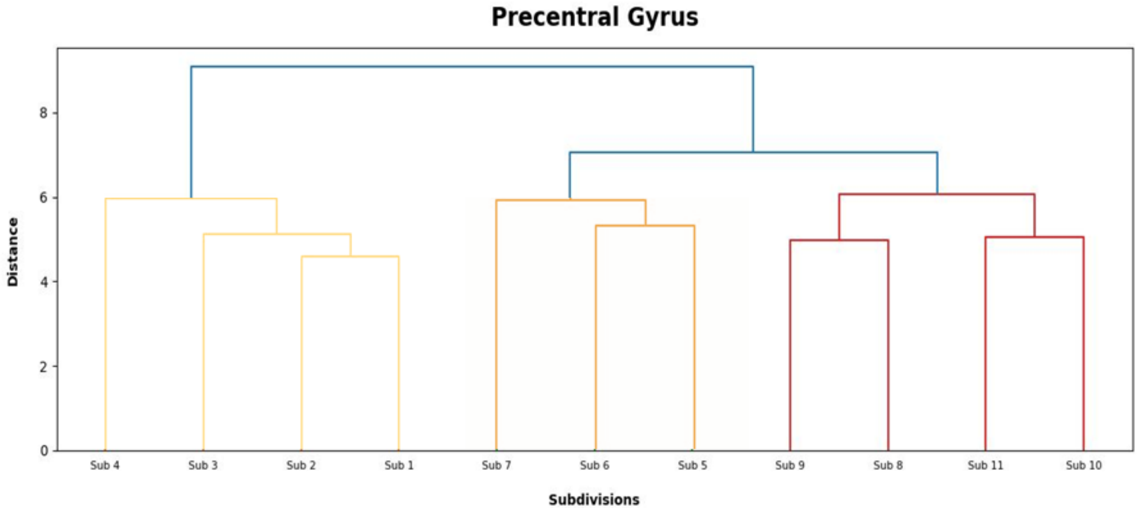

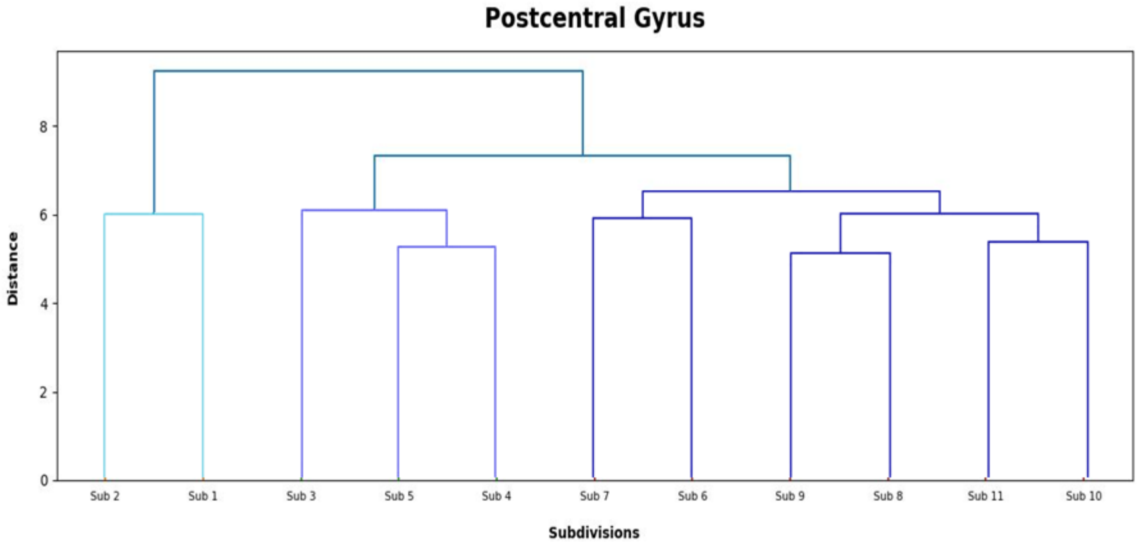


**Supp. Figure 2.** Dendrograms derived from agglomerative hierarchical clustering. Each color represents an individual cluster in pre-central gyrus (top figure): $Pre_{LAT}$, $Pre_{INT}$, $Pre_{MED}$ (yellow, orange, red, respectively); and post-central gyrus (bottom figure): $Post_{LAT}$, $Post_{INT}$, $Post_{MED}$ (light blue, blue, indigo, respectively).

**Supp. Table 1.** Comparisons of $FT_{Ref}$ and $FT_{Ind}$ groups per cluster: t-tests for each diffusion metric. P-values are corrected for multiple comparisons. Significant results are highlighted in green. See Table 1 for subject group legend and Figure 1 for cluster legend.

**$FT_{Ref}$ vs. $FT_{Ind}$**

| | AD | | RD | | MD | | FA | | NDI | | ODI | |
|---|---|---|---|---|---|---|---|---|---|---|---|---|
| | T | $p_{corr}$ | T | $p_{corr}$ | T | $p_{corr}$ | T | $p_{corr}$ | T | $p_{corr}$ | T | $p_{corr}$ |
| $Pre_{LAT}$ | -1.78 | 0.274 | -1.53 | 0.341 | -1.72 | 0.274 | -0.33 | 0.811 | 1.74 | 0.274 | -2.29 | 0.159 |
| $Pre_{INT}$ | -0.80 | 0.570 | -0.53 | 0.716 | -0.58 | 0.698 | -0.12 | 0.934 | 0.45 | 0.735 | 4.16 | 0.001 |
| $Pre_{MED}$ | -1.20 | 0.550 | -0.85 | 0.570 | -0.86 | 0.570 | -0.19 | 0.900 | 0.73 | 0.603 | 5.02 | <0.001 |
| $Post_{LAT}$ | -1.64 | 0.298 | -1.97 | 0.245 | -1.89 | 0.257 | 0.80 | 0.570 | 2.13 | 0.194 | -6.74 | <0.001 |
| $Post_{INT}$ | -1.06 | 0.550 | -0.99 | 0.550 | -1.08 | 0.550 | 0.49 | 0.731 | 1.11 | 0.550 | -12.06 | <0.001 |
| $Post_{MED}$ | -1.02 | 0.550 | -0.94 | 0.550 | -0.96 | 0.550 | 0.02 | 0.988 | 0.97 | 0.550 | -8.56 | <0.001 |

**Supp. Table 2: Comparison between $FT_{Ref}$ and $PT_{TEA}$ groups in terms of dMRI metrics. a,** Results of ANCOVA analysis examining the relationship between each diffusion metric and covariates, including PMA at scan, residual metric over whole cortex (corrected for PMA at scan), and factors of interest: group ($FT_{Ref}$, $PT_{TEA}$), region (pre-/post-central gyrus), and position (lateral / intermediary / medial), along with relevant interactions. The models account for hierarchical structure of the repeated measures across matched subjects, regions, and positions, with a nested error term. *Legend:* F: F-statistic from ANCOVA; p: statistical significance; partial $\eta^2$ = effect size. **b,** Post-hoc group comparison: paired t-tests between $FT_{Ref}$ and $PT_{TEA}$. P-values are corrected for multiple comparisons. Significant results are highlighted in green. See Table 1 for subject group legend and Figure 1 for cluster legend.

**a,**

| | **AD** | | | **RD** | | |
|---|---|---|---|---|---|---|
| | **F** | **p** | **Partial η²** | **F** | **p** | **Partial n2** |
| ***Covariates*** | | | | | | |
| Residual GM | - | - | - | - | - | - |
| PMA at scan | 107.44 | <0.001 | 0.09708 | 88.30 | <0.001 | 0.08387 |
| ***Factors*** | | | | | | |
| Group | 179.98 | <0.001 | 0.16261 | 172.98 | <0.001 | 0.16431 |
| Region | 12.48 | <0.001 | 0.01128 | 11.43 | 0.001 | 0.01085 |
| Position | 55.08 | <0.001 | 0.09953 | 42.31 | <0.001 | 0.08037 |
| ***Interactions*** | | | | | | |
| Group:Region | 2.06 | 0.152 | 0.00186 | 1.14 | 0.285 | 0.00109 |
| Group:Position | 0.84 | 0.433 | 0.00151 | 0.67 | 0.514 | 0.00127 |
| | **MD** | | | **FA** | | |
| | **F** | **p** | **Partial η²** | **F** | **p** | **Partial n2** |
| ***Covariates*** | | | | | | |
| Residual GM | - | - | - | 726.75 | <0.001 | 0.47415 |
| PMA at scan | 95.62 | <0.001 | 0.08875 | 2.10 | 0.147 | 0.00137 |
| ***Factors*** | | | | | | |
| Group | 179.31 | <0.001 | 0.16642 | 62.82 | <0.001 | 0.04099 |
| Region | 13.10 | <0.001 | 0.01216 | 4.95 | 0.026 | 0.00323 |
| Position | 46.90 | <0.001 | 0.08705 | 15.25 | 0.000 | 0.01990 |
| ***Interactions*** | | | | | | |
| Group:Region | 1.24 | 0.266 | 0.00115 | 0.83 | 0.363 | 0.00054 |
| Group:Position | 0.69 | 0.500 | 0.00129 | 6.39 | 0.002 | 0.00834 |
| | **NDI** | | | **ODI** | | |
| | **F** | **p** | **Partial η²** | **F** | **p** | **Partial n2** |
| ***Covariates*** | | | | | | |
| Residual GM | 2163.11 | <0.001 | 0.50951 | 247.35 | <0.001 | 0.18895 |
| PMA at scan | 295.79 | <0.001 | 0.06967 | 141.36 | <0.001 | 0.10798 |
| ***Factors*** | | | | | | |
| Group | 703.64 | <0.001 | 0.16574 | 10.95 | 0.001 | 0.00837 |
| Region | 47.16 | <0.001 | 0.01111 | 80.69 | <0.001 | 0.06164 |
| Position | 167.19 | <0.001 | 0.07876 | 53.67 | <0.001 | 0.08200 |
| ***Interactions*** | | | | | | |
| Group:Region | 5.58 | 0.018 | 0.00131 | 26.60 | <0.001 | 0.02032 |
| Group:Position | 1.89 | 0.152 | 0.00089 | 1.40 | 0.248 | 0.00214 |

**b,**

### $FT_{Ref}$ vs. $PT_{TEA}$

| | AD | | RD | | MD | | FA | | NDI | | ODI | |
|---|---|---|---|---|---|---|---|---|---|---|---|---|
| | T | $p_{corr}$ | T | $p_{corr}$ | T | $p_{corr}$ | T | $p_{corr}$ | T | $p_{corr}$ | T | $p_{corr}$ |
| **$Pre_{LAT}$** | -5.58 | <0.001 | -5.04 | <0.001 | -5.29 | <0.001 | 1.32 | 0.199 | 5.21 | <0.001 | 1.58 | 0.132 |
| **$Pre_{INT}$** | -4.12 | <0.001 | -4.24 | <0.001 | -4.28 | <0.001 | 2.62 | 0.014 | 4.24 | <0.001 | -2.29 | 0.032 |
| **$Pre_{MED}$** | -4.26 | <0.001 | -4.3 | <0.001 | -4.32 | <0.001 | 3.16 | 0.003 | 4.29 | <0.001 | -1.40 | 0.178 |
| **$Post_{LAT}$** | -5.67 | <0.001 | -5.6 | <0.001 | -5.62 | <0.001 | 1.94 | 0.065 | 5.55 | <0.001 | 2.13 | 0.044 |
| **$Post_{INT}$** | -5.41 | <0.001 | -5.09 | <0.001 | -5.20 | <0.001 | 0.81 | 0.424 | 5.08 | <0.001 | 4.08 | <0.001 |
| **$Post_{MED}$** | -5.17 | <0.001 | -5.28 | <0.001 | -5.31 | <0.001 | 2.96 | 0.006 | 5.31 | <0.001 | 2.16 | 0.042 |

**Supp. Table 3**. Assessment of the nullity of Mahalanobis distances in $PT_{Birth}$ and $PT_{TEA}$ groups: t-tests for each cluster are compared to 0 (i.e. mean of the reference group). P-values are corrected for multiple comparisons. See Table 1 for subject group legend and Figure 1 for cluster legend.

| | $PT_{Birth}$ | | $PT_{TEA}$ | |
|---|---|---|---|---|
| | T | $p_{corr}$ | T | $p_{corr}$ |
| **$Pre_{LAT}$** | 38.516 | <0.001 | 24.876 | <0.001 |
| **$Pre_{INT}$** | 45.321 | <0.001 | 26.929 | <0.001 |
| **$Pre_{MED}$** | 41.567 | <0.001 | 19.733 | <0.001 |
| **$Post_{LAT}$** | 32.000 | <0.001 | 23.703 | <0.001 |
| **$Post_{INT}$** | 40.994 | <0.001 | 27.102 | <0.001 |
| **$Post_{MED}$** | 40.994 | <0.001 | 22.937 | <0.001 |
